# Personalized smartphone notifications bias auditory salience across processing stages

**DOI:** 10.64898/2026.01.26.701556

**Authors:** Prakash Mishra, Tapan K. Gandhi, Saurabh R. Gandhi

## Abstract

Modern digital environments expose the brain to a dense stream of personally meaningful cues that differ markedly from the conditions under which sensory systems evolved. Understanding how the brain responds and adapts to such environments is increasingly relevant for cognitive and mental well-being. In this study, we used auditory event-related potentials (ERPs) recorded during a task-irrelevant auditory oddball paradigm to obtain a time-resolved account of neural responses to smartphone notifications. We dissociated the effects of generic notification sounds from the learning-related effects of personalized notification sounds, and examined how these responses varied across levels of smartphone use. Among early stages of processing, generic tones exhibited experience-dependent strengthening of auditory representations, whereas personalized tones engaged stable, pre-learned representations, as reflected in P2 dynamics. The strongest stimulus-dependent modulation was observed in pre-attentive salience detection, reflected in the mismatch negativity (MMN), which was earlier and larger for personalized tones compared to generic ones. At later stages, attentional orienting and extended evaluative processing showed distinct patterns that were differentially sensitive to stimulus identity and smartphone use. Notably, smartphone usage levels modulated later attentional and evaluative stages (P3a and LPP) without altering early pre-attentive prediction-error signaling. Overall, our findings demonstrate that personalized smartphone notifications do not uniformly increase neural distraction. Instead, they engage robust and rapid pre-attentive prediction-error signaling that is largely independent of usage, while habitual smartphone use selectively shapes later attentional orienting and evaluative dynamics.

**Significance statement:** Modern digital environments are saturated with personalized cues, such as smartphone notifications, that are known to disrupt ongoing behavior. A common assumption in discussions of digital distraction is that repeated exposure to such cues progressively amplifies their sensory salience, rendering habitual users increasingly reactive. Here, we challenge this assumption using an ecologically grounded neural paradigm with personalized notification sounds. We show that personalized notifications engage robust early salience detection that is shared across users, while habitual smartphone use selectively modulates later attentional and evaluative processing. These findings place important constraints on how repeated exposure to personalized digital cues shapes neural processing, indicating that usage-related effects emerge downstream rather than at early sensory stages.

## Introduction

Smartphone notifications have been shown to cause distraction, anxiety and performance deterioration through several behavioral studies ^1–5^. For example, people respond more slowly and with greater mental effort when hearing a phone notification, even if they are focused on an unrelated task ^6,7^. Importantly, smartphone alerts are highly personalized auditory signals. Users select custom ringtones or app-specific chimes that carry personal or social meaning (e.g. a friend’s message or a social media ping). These personalized signals often recruit higher order cognitive processes involving emotions, rewards and social motivations, even giving rise to a sense of *“fear of missing out”* ^8,9^. In fact, individuals more susceptible to such fear have shown a tendency to stop their current activities more often when they receive an interruptive notification ^10^. These studies substantiate the detrimental behavioral impact of notification sounds specifically personalized to the user.

Behavioral metrics capture only the final outcome of complex perceptual, evaluative, and motivational processes, and are often insensitive to early, implicit, or cumulative effects ^4,11^. Conscious and unconscious neural processes operating over slower timescales may accumulate without being reflected in immediate behavior. For example, craving-related responses can be actively suppressed ^12^, and subtle shifts in attention may not be detectable at the behavioral level ^13^. Even when individual notifications do not elicit overt responses, repeated exposure may gradually bias perceptual and attentional systems ^14^, increasing the likelihood of future orienting, distraction, or checking behavior. Neural measures provide access to these missing intermediate computational stages that link stimulus exposure to behavioral outcomes. Therefore, assessing the neural impact of smartphone notifications is essential for developing a nuanced, mechanistic understanding of how they influence behavior.

Surprisingly, however, the neural impact of notifications is relatively underexplored. In the pre-smartphone era, Event Related Potentials (**ERPs**) evoked by personalized notifications have been shown to have an altered mismatch negativity (**MMN**), including a late negative deflection and an enhanced late P3a ^15,16^. These boosted responses indicate that self-relevant sounds automatically draw attention, suggesting that personal significance shapes auditory processing through long-term memory effects. With the explosive use and misuse of smartphones since, personalized notifications are expected to have stronger neural signatures. A recent EEG study assessing the neural impact of notifications during a visual Go/No-Go task found them to disrupt cognitive processing, reflected in reduced N200 and P300 amplitudes. The effects were more severe and prolonged in individuals at risk of overuse ^17^. The non-risk group also exhibited higher error rates and impulsive responses, indicating greater vulnerability to distraction. However, the study used artificial vibratory stimuli applied behind the participants as a proxy for notifications, potentially confounding higher order effects with generic distractor effects. Thus, ERP (or any neural) studies of the impact of smartphone notifications, especially their personalized nature, are currently lacking.

Auditory ERP responses can provide a detailed account of the neural processes recruited by personalized notification sounds at a high temporal resolution, spanning across the pre-attentive and attention-related processing stages ^18–20^ (Fig. 1A). Within the broader umbrella of late-latency auditory evoked potentials starting with P1 onwards, early responses such as N1 and P2 can reflect heightened neural sensitivity to certain sounds, driven by increased sensory gain modulated by top-down expectation, and learned associations respectively ^21–23^. Mid latency responses such as the MMN reflect the pre-attentive responses that may be modulated based on the salience of the sound, associated with automatic deviance detection and closely coupled to orienting responses that engage autonomic systems ^24–26^. Lastly, late responses such as the P3a and late positive potential (LPP) can reflect attention shifts and emotional salience respectively ^27,28^. Therefore, in the present study, we compare each of the above auditory ERP responses across personalized and generic auditory stimuli and across a spectrum of smartphone users to build a time-resolved and process-specific understanding of the impact of personalized notification sounds.

**Figure 1.**
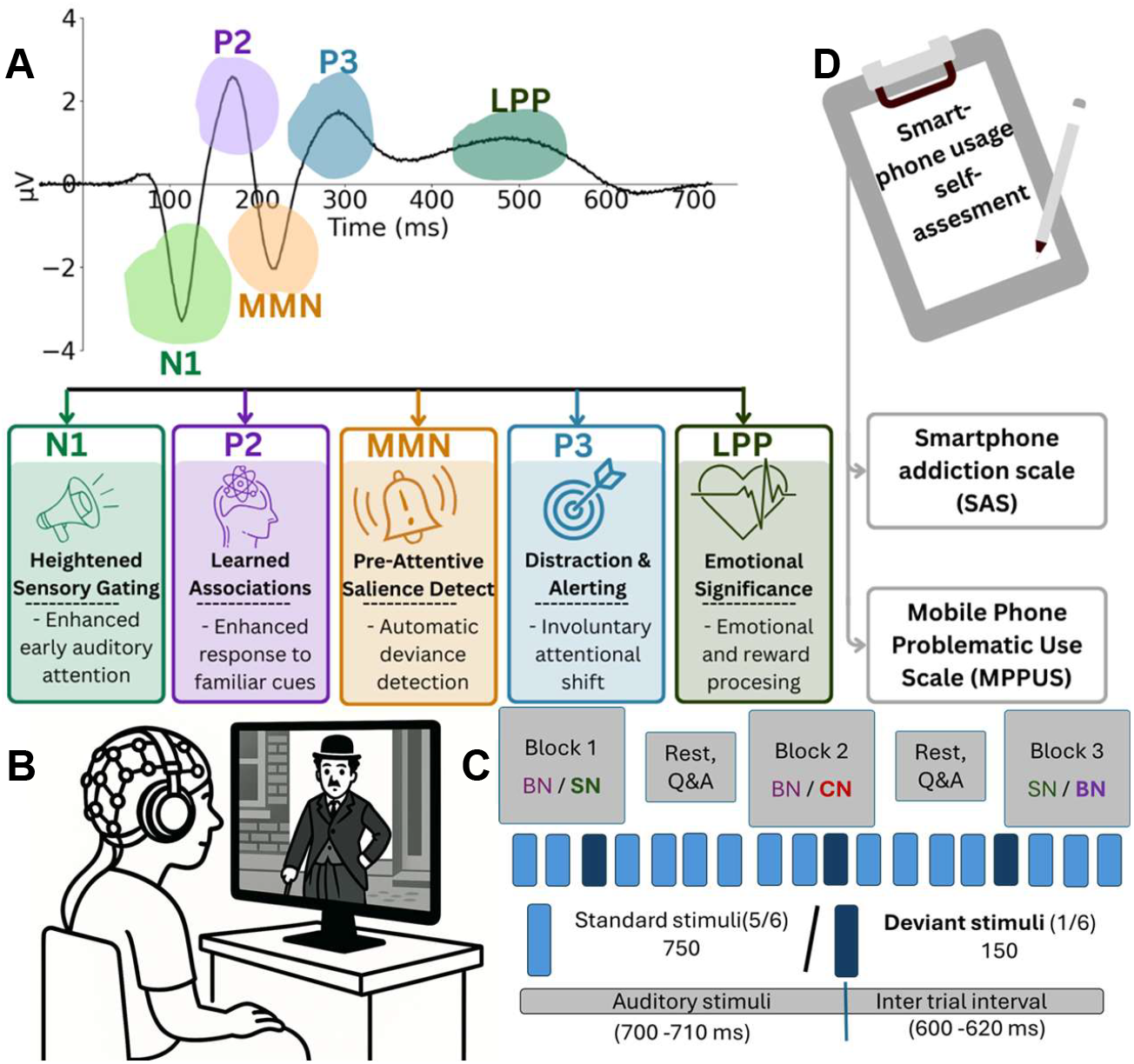
Experiment design and hypotheses. **(A)** Schematic event-related potential (ERP) waveforms illustrating key auditory components, N1, P2, mismatch negativity (MMN), P3, and late positive potential (LPP), with their approximate time windows and associated cognitive-emotional processes. **(B)** Behavioral self-report measures used in the study: smartphone usage self-assessment, the Smartphone Addiction Scale (SAS), and the Mobile Phone Problematic Use Scale (MPPUS). **(C)** Schematic of the EEG recording setup, where participants were exposed to three auditory stimuli via headphones, Smartphone Notification (SN), Control Notification (CN), and Beep Notification (BN), while viewing a silent movie. **(D)** Task structure showing stimulus blocks, rest/Q&A periods, and the timing of standard and deviant auditory stimuli with inter-trial intervals.

We hypothesize that smartphone notifications can recruit distinct neural processes based on an individual’s level of exposure and use. Because personal ringtones are highly salient, we expect them to evoke faster pre-attentive responses (MMNs) compared to generic tones. We also anticipate that users who heavily engage with their phones will show different neural signatures than light users, although the specific effects may not be obvious. For example, overlearning may attenuate novelty-driven orienting (reduced P3a), whereas heightened salience may enhance attentional capture. The present study adjudicates between these alternatives. Because these tones carry emotional and motivational value, HSUs might exhibit an amplified LPP in response to specific personalized notifications, indicating greater evaluative or emotional processing. In contrast, light smartphone users (**LSU**s) may find the notification less familiar and less emotionally loaded, yielding the opposite patterns. Therefore, in addition to building a time-resolved understanding of cognitive processes evoked by personalized notifications, we also investigate the dependence of these processes on the level of smartphone use.

In order to assess the entire hierarchy of auditory ERPs in response to notification sounds, we used a task-irrelevant auditory oddball paradigm with personalized and control notification sounds (Fig. 1B). Although the key feature of our study is the use of personalized notifications, applying ERP methods across the wide variety of notification sounds presents notable methodological difficulties. Notification sounds differ markedly in pitch, timbre, duration, and loudness (Fig. S1), making it challenging to compute a conventional MMN by subtracting a single ‘standard’ tone from each participant’s notification deviant. We therefore adopted an across-block difference wave design, where the standard and deviant are interchanged between the pair of blocks (Fig. 1C). Personalized notification sounds were compared against a standard beep sound as well as a control extended notification sound that the subjects were unfamiliar with. Finally, to probe usage-dependent modulation, all participants completed a self-reported behavioral questionnaire for problematic smartphone use and addiction (Fig. 1D). Our experiment design allows us to use raw as well as appropriate difference waves to test hypotheses regarding the different ERP components and their modulation by usage intensity.

## Methods

### Participants

A total of 57 (male = 43, female = 14) participants’ EEG data were collected. Out of these, two subjects were removed due to trigger marking errors. 43 participants were college-going students aged 18 to 31 years. 14 participants were housekeeping staff from a low-education and low-income background, aged between 20 and 48 years.

The experiment design was approved by the IIT Delhi Institutional Ethics Committee (approval number 2021/P0143). All participants were provided with detailed information about the protocol and potential hazards, and filled out a consent form accordingly before initiating the experiment.

### Experiment protocol

The experiment protocol was based on the task-irrelevant auditory oddball paradigm (Fig. 1B,C). Three types of auditory stimuli were provided: the beep notification tone (**BN**), the personal smartphone notification tone (**SN**), used by the respective participants, and a control notification tone (**CN**), which was also a smartphone notification, but unfamiliar to the participants, used uniformly across a subset of participants. SN served as the personally significant tone, BN served as a non-significant baseline, while CN, with its extended temporal structure, served as a non-significant control.

The experiment consisted of two primary blocks, along with one additional block for 25 participants, which was counterbalanced with the primary blocks. In the two primary blocks, SN and BN served as the standard (deviant) and deviant (standard) respectively. In the third block, BN served as the standard and CN as the deviant, to control for the extended nature of SN which was missing in BN. This also allows direct comparison to the previous literature ^29^. The order of these blocks was counterbalanced across participants to control for presentation order bias.

The experiment lasted for 60 mins, with 20 mins for each of the three blocks. In each block, a total of 900 auditory stimuli were provided, with ⅙ (150) as deviants and ⅚ (750) as standards. The number of standards between two consecutive deviants varied uniformly between three and nine. Each trial began with an auditory stimulus lasting between 700–710 ms to reduce predictability. Longer tones were clipped at the length of the stimulus period and shorter tones were left unchanged. This was followed by an inter-trial interval of 600–610 ms. The total duration of each trial was thus approximately 1300 ms.

The participants watched a silent, but engaging black-and-white movie, *Charlie Chaplin’s Modern Times* (short clips were adopted for use under fair use principles), while sitting in a comfortable chair, and received auditory stimuli from on-ear headphones. Participants were explicitly instructed to ignore the auditory stimuli and focus on the video at the start of the experiment. To assess participants’ attentiveness to the video during the task, we posed five questions related to the storyline after each block. The movie was displayed sequentially without disrupting the storyline irrespective of the ordering of the blocks.

The experiment protocol was designed and presented using the PsychoPy-2024.1.4 toolbox with Python version 3.8.10 on a Windows 11 computer, on a 27 inch screen placed 120 cm away from the participants’ face. Participants were not head-fixed, though instructed to minimize head movements, including laughter.

Self-reported behavioral metadata relating to phone usage was collected at the end of the experiment via a Google form.

### Data collection

#### EEG data

We collected electroencephalogram (EEG) data during the task using the 64-channel actiCHamp system. Electrodes were placed according to the 10-20 system with the Cz electrode acting as reference while recording. Data was collected at a sampling rate of 1000 Hz. Heart rate and skin conductance sensor data was acquired through the actiCHamp’s auxiliary channels for participants who completed the CN block.

#### Behavioral metadata

Following EEG data collection during the task, we gathered participants’ responses on the Smartphone Addiction Scale (SAS) (Li et al., 2023) and Mobile Phone Problematic Use Scale (MPPUS) (Mach et al., 2020) questionnaires along with additional smartphone use related self-reported data (see SI).

### EEG preprocessing

EEG data preprocessing and analysis were performed using the MNE-Python package (version 1.7.0) and Python (version 3.12.3). Raw EEG signals were re-referenced to the average of the mastoid electrodes (TP9 and TP10) and subsequently downsampled to 512 Hz. A bandpass filter between 0.1 to 30 Hz was then applied to the data. Independent component analysis (ICA) was then conducted to identify and remove ocular artifacts. Ocular artifacts were detected and removed using MNE’s automated electrooculogram (EOG) component identification algorithm, based on the correlation between component and frontal channels (Fp1, Fp2, AF7, AF8, F7 and F8) activity.

EEG epochs were segmented from -300 ms to 900 ms relative to stimulus onset, with a baseline correction applied using the 300 ms pre-stimulus interval. Any channel exhibiting a peak-to-peak voltage exceeding 80 µV within an epoch was flagged as a bad channel. If more than 10% of channels (i.e. more than six electrodes) were marked as bad in a given epoch, that epoch was excluded from further analysis. Otherwise, the identified bad channels were corrected using spline interpolation. Finally, a visual inspection was performed to identify and remove epochs showing excessive muscle or movement artifacts.

### Data analysis

#### Standardized Measurement Error (SME) – based ERP metric computation

To assess the reliability of ERP measures and identify low-quality data, standardized measure error (**SME**) was computed for each participant, while computing peak latency and the time-window mean amplitude ^30^. SME based ERP measures were computed by bootstrapping over trials in each block and condition to obtain stable estimates of amplitude and latency.

For each participant, block and condition, epochs were resampled 500 times with replacement and averaged to generate a bootstrapped ERP waveform *E*(*t*). Peak latency was extracted within a predefined window [*t*_l_, *t*_2_] as *PL* = *arg max* (± *E*(*t*)); *t* ∈ [*t*_l_, *t*_2_]; depending on the polarity of the component. To obtain the mean amplitude, the waveform was first detrended by removing the linear baseline connecting the start and end of the analysis window, ensuring that the area-based measure reflected true ERP morphology rather than slow drift. The detrended waveform *E*_*d*_(*t*) was integrated to obtain the window area *A* = Σ *E*_*d*_(*t*), from which mean amplitude was computed as *A*/(*t*_2_ − *t*_l_). For difference-wave analyses, the ERPs from the two conditions were averaged separately and subtracted as *D*(*t*) = *E*_0_(*t*) − *E*_l_(*t*) prior to scoring in each bootstrap iteration. Repeating this procedure across iterations yielded bootstrap distributions of peak latency and mean amplitude, providing robust parameter estimates.

#### Time window selection and statistical tests for ERP analysis

The time windows of interest for analysis of late components (P3a and LPP) were identified using a cluster-based permutation test (5,000 permutations) applied to the grand-average ERPs, with a cluster-forming threshold of p = 0.05. Only clusters consistent with the canonical latency range of the ERP component under investigation were considered, and the resulting cluster-derived time windows were used for metric extraction and statistical evaluation.

Significance of within-subject comparison across tones was assessed using paired t-tests. Significance of between-group comparisons was assessed using unpaired t-tests.

#### N1 alignment of ERPs

Unlike simple tonal stimuli, smartphone notification sounds are acoustically complex and do not exhibit an immediate salient onset. Many notification sounds contain a delayed rise to the acoustic event that triggers perceptual detection (Fig. S1), making it challenging to align and compare ERPs across distinct SN stimuli. However, the N1 ERP component is known to align with perceptual rather than physical sound onset ^31^. We therefore time-shift all ERPs to align with the N1 peak of responses evoked by the standard tones, thereby setting up a common temporal baseline across subjects (SN tones) for subsequent analysis. Aligning ERP responses to their tone-specific standard N1 peaks minimizes acoustic onset confounds and ensures that the resulting difference waves more accurately reflect underlying neurocognitive processing rather than acoustic differences.

Further, a within-block analysis was performed for all subjects who completed the control block. Since the CN tone is never used as a standard, for this analysis, the standard and deviant tones from each block were aligned to their own respective N1 peaks prior to computing the difference waves. The contrasts CN-BN, SN-BN, and BN-SN were then obtained by subtracting the STD N1-aligned standard response of each tone from its corresponding DEV N1-aligned deviant response within that block.

#### Spectral clustering to classify heavy and light smartphone users

To identify different smartphone user groups, a spectral clustering machine learning algorithm was applied to the SAS responses of the participants. Instead of using participants’ total SAS scores, responses to individual questionnaire items were used to capture detailed response patterns. All items were standardized to ensure equal contribution of each question. A similarity structure among participants was then constructed based on their response patterns, and spectral clustering was used to group participants according to this structure. The resulting clusters were derived from a low-dimensional representation of the data and partitioned using k-means clustering. These clusters were interpreted as heavy, average and light smartphone-use groups based on their SAS response patterns. Spectral clustering was implemented in Python using the scikit-learn (version 1.5.0) library; a mathematical description is provided in the supplementary materials.

## Results

### N1 – P2 SEM based subject selection and ERP alignment

Given the non-uniformity of SN stimuli across subjects, we used the N1 peak as a uniform marker for acoustic perceptual onset, aligning all ERPs to this timepoint for further comparison. This choice is substantiated by the fact that the observed variation in N1 latency across subjects with the same SN tone is comparable to the variation in N1 latency of the BN tone, which is uniform across all participants (Fig. S2).

To clearly distinguish the MMN from the attention-related auditory N2pc component in later analysis, we restricted the MMN window between the N1 and P2 peaks for each subject respectively. Consequently, we required subjects to display robust and reliable N1 and P2 for inclusion in further analysis. A total of 42 subjects showed both N1 and P2 peaks reliably, with peak latencies for the standard stimulus having standard deviations of up to a maximum of 0.016 ms (N1) and 0.0125 ms (P2), respectively after bootstrapping (Fig. S3), and were retained. The remaining subjects were dropped from the analysis since the acoustic onsets cannot be determined reliably without a reliable N1, nor the MMN window without a reliable P2.

For each subject, the N1 peak latency for the standard tone was first computed by bootstrapping over trials and taking the mean peak latency across bootstrapping iterations. All epochs were then time-shifted to align with the corresponding standard-stimulus N1 peak latency (ie SN STD and DEV both were aligned to SN STD N1, and similarly for BN tones) for further analyses.

### Tone-dependent modulation for selective early auditory ERP components

Repetition-related modulation was evident across both early and later stages of auditory processing. For standard stimuli, pronounced attenuation was observed in the early negative time window corresponding to the N1 component. N1 mean amplitude decreased significantly from the first to the last trial quartile for both SN (Q1: −2.04 µV; Q4: −1.64 µV; *t*(54) = −5.58, *p* <.001; Fig. 2A) and BN tones (Q1: −2.09 µV; Q4: −1.48 µV; *t*(54) = −6.75, *p* <.001; Fig. 2C), indicating robust early sensory adaptation. In the subsequent P2 time window, visual inspection of the grand-average ERPs did not reveal an obvious change from Q1 to Q4 for either stimulus type (Fig. 2A and C).

**Figure 2.**
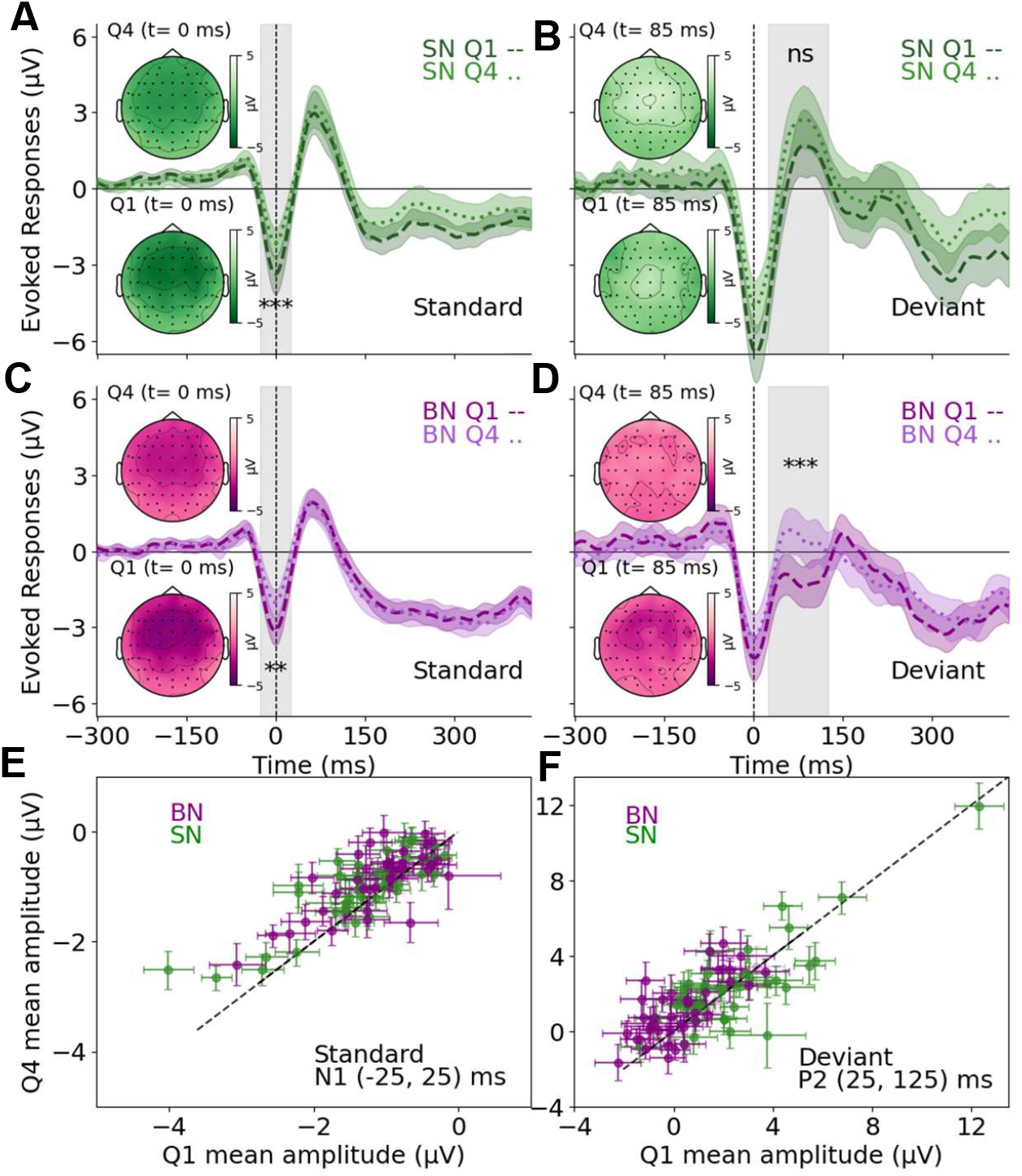
Adaptation effects in N1 and P2 for smartphone (SN) and beep notifications (BN). Grand-average ERP waveforms for deviant stimuli and corresponding scalp topographies are shown for the first (Q1; dashed lines) and last (Q4; dotted lines) quartiles of trials. **A, B**. Responses to SN tones (green). **C, D**. Responses to BN tones (purple). Shaded regions represent 95% confidence interval across participants. Scalp maps illustrate the spatial distribution of activity in the N1 (left column; negative) and P2 (right column; positive) time windows. **E, F**. Mean N1 amplitudes for SN and BN stimuli as STD and DEV in quartile 4 vs quartile 1 for individual subjects. Error bars represent SEM.

For deviant stimuli, N1 amplitudes did not show significant modulation across quartiles for either stimulus type (SN: Q1: −2.91 µV, Q4: −2.65 µV; *t*(41) = −1.31, *p* =.20; BN: Q1: −2.18 µV, Q4: −2.17 µV; *t*(41) = −0.07, *p* =.95). However, clear stimulus-dependent differences emerged in the P2 time window. While SN deviants showed stable P2 amplitudes across repetitions (Q1: 2.28 µV; Q4: 2.36 µV; *t*(41) = −0.34, *p* =.74; Fig. 2B), BN deviants exhibited a significant increase in P2 amplitude from Q1 (0.38 µV) to Q4 (1.18 µV; *t*(41) = −4.49, *p* <.001; Fig. 2D), consistent with repetition-related learning effects.

The large number of trials in our experiment design allowed us to perform individual-level analysis that further corroborated the population level observations. For standard stimuli, N1 mean amplitudes in the first quartile were consistently more negative than those in the final quartile for a majority of individuals for both SN (32/42) and BN (31/42) (Fig. 2E), demonstrating clear repetition-related adaptation of early auditory processing. On the other hand, for deviant P2 responses, we found a much more even distribution of increase vs decrease of amplitude between the first and last quartile, slightly favouring an increase (26/42 subjects showed a significant increase). For BN P2 deviants, again, a majority of the subjects (32/42) exhibited larger mean amplitudes in the fourth quartile than in the first quartile (Fig. 2F). This pattern was consistent with repetition-related learning or strengthening of auditory representations for BN, which the participants are not familiar with, compared to personalized SN, which is presumably overlearned.

### Increased amplitude and reduced latency of auditory MMNs for personalized notification tones

For analyzing the MMN, we adopted an across-block difference wave design (for example, SN MMN = SN DEV from block 1 - SN STD from block 2) to mitigate ERP artifacts due to the extended nature of smartphone notifications compared to beeps. We looked for the MMN in the window between the N1 and P2 peaks to avoid confounds due to the attention-related N2pc. For the grand averaged difference waves, we defined this window from t = 0 (N1 peak, common across subjects after alignment) to t = 70 ms, the average P2 latency across subjects (Fig. 3A). We clearly observed the MMN ERP in this period with a consistent fronto-central scalp distribution for both SN and BN (Fig. 3B, inset). Moreover, the grand averaged SN MMN at the fronto-central electrodes (Fz, FCz, and Cz) was significantly larger and earlier compared to BN (Fig. 3B).

**Figure 3.**
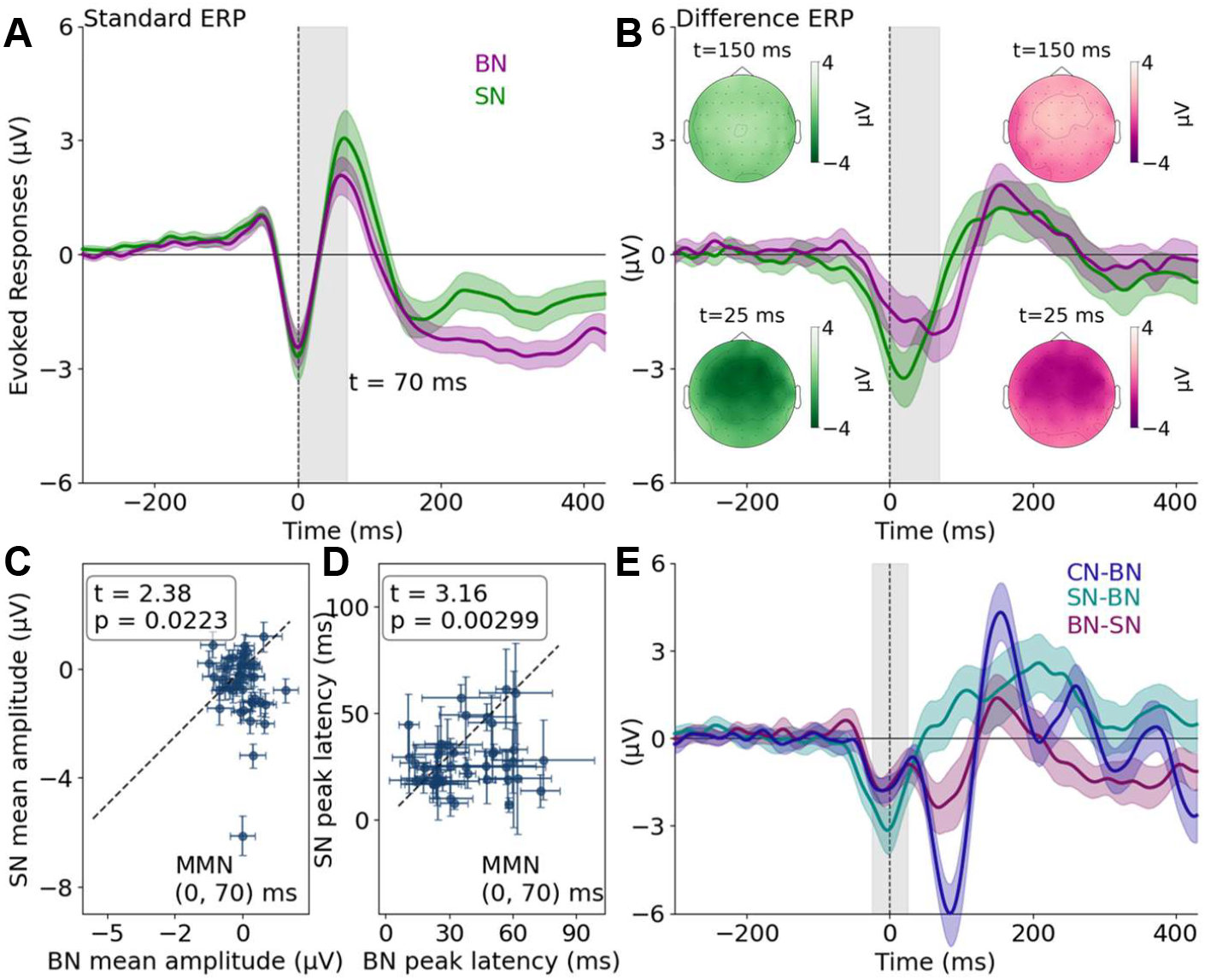
Mismatch negativity (MMN) response for smartphone (SN), beep (BN), and control notifications (CN). (A) Grand-averaged ERPs time-aligned to each participant’s N1 peak for the standard tone in BN (purple) and SN (green) conditions. (B) MMN waveforms computed by subtracting standard-tone ERPs from deviant-tone ERPs, time-locked to the standard-tone N1 peak. (Insets) Scalp topographies (25 ms post-MMN onset) for SN (green) and BN (purple). (C) SN vs BN mean MMN amplitudes for individual subjects. (D) SN vs BN peak MMN latencies for individual subjects. Each point in D, E reflects a single participant, with error bars showing within-subject standard deviation. (E) Within-block MMNs: SN (DEV) – BN (STD) (teal), BN (DEV) – SN (STD) (magenta) and CN (DEV) – BN (STD) (blue). A, B, E: Shaded areas represent 95% confidence intervals; the time window marks the N1– P2 interval used for ERP metric extraction.

For a within-subject analysis, we restricted the MMN to the window between the respective N1 and P2 peaks. Within these windows, we observed that the SN MMN amplitudes were consistently lower than BN MMN for a majority of subjects, although variations were seen with a few subjects showing a significantly lower SN MMN amplitude compared to BN MMN (Fig. 3C). The within-subject latency comparison was even stronger: only a small minority of subjects (13/42) showed a marginally longer latency for SN MMN, while most subjects showed a substantially shorter SN MMN latency (Fig. 3D).

These results clearly reflect a stable difference in pre-attentive processing of personally significant and non-significant sounds with variations across SN tones accounted for. However, SN and BN tones differ substantially in that BN has no temporal modulations compared to SN tones. To control for this, we performed a within-block analysis, by computing MMNs for the CN, SN and BN DEV after subtracting the standard from the same block. Since the STD and DEV are distinct tones, for this analysis we aligned the ERPs with the respective N1 peaks (Fig. S4). We note that for within-block difference waves, only very early modulations immediately surrounding the N1 peak (auditory stimulus onset) are meaningful. At later times, the difference waves are confounded by the differences in the temporal modulations of the STD and DEV tones.

In the within-block difference waves, we observed an MMN-like negative deflection close to 0, although peaking at slightly negative times (Fig. 3E), likely an artifact of differential alignment of the individual ERPs to their respective N1 peaks instead of a common alignment (Fig. S5). Yet, the SN MMN deflects downwards earlier compared to BN and CN, which are virtually identical. Moreover, SN MMN amplitude was significantly larger than CN and BN, both of which are identical. These observations further reaffirm the increased amplitude and reduced latency of SN MMNs compared to other sounds, irrespective of the temporal structure or overall category of the sound.

### Comparable P3a but reduced LPP amplitude for SN at later times

In the later positive time window corresponding to the novelty P3 (P3a) component (140.625–171.875 ms), the BN condition showed a larger positive peak than the SN condition (Fig. 3B), although the peak latency did not differ between conditions (SN: 156.4 ± 0.0043 ms; BN: 156.2 ± 0.0043 ms; t = 0.18, p = 0.854). The mean amplitude was larger for BN (0.152 ± 0.268 μV) compared to SN (0.064 ± 0.269 μV), but again this difference did not reach statistical significance (t = -1.17, p = 0.249). Together, these findings indicate that although the BN condition elicited a visually stronger P3a response, the amplitude and timing differences were not statistically reliable at the group level. However, in the subsequent window corresponding to the late positive potential (LPP) (310–360 ms), the mean amplitude was more negative for SN (-0.137 ± 0.327 μV) compared to BN (0.009 ± 0.318 μV) with statistical significance (t = -2.79, p = 0.017) at parieto-central (Pz, CPz, Cz) electrodes. Peak latency during this window did not differ significantly (t = -0.41, p = 0.687).

### Smartphone usage modulates ERP characteristics

#### Participant Stratification and Psychometric Validity

Participants were subsequently grouped into the Heavy Smartphone Users (HSU), Light Smartphone Users (LSU), and Average Smartphone Use (ASU) categories based on their SAS scores. The psychometric reliability of both self-report scales was high, with Cronbach’s α = 0.903 for the SAS and α = 0.854 for the MPPUS. Shapiro–Wilk tests indicated that the distributions of both SAS (W = 0.978) and MPPUS (W = 0.983) scores did not substantially deviate from normality. The two measures were strongly correlated (r = 0.756, p = 0), demonstrating convergent validity across independent assessments of smartphone use (Fig. 4C).

**Figure 4.**
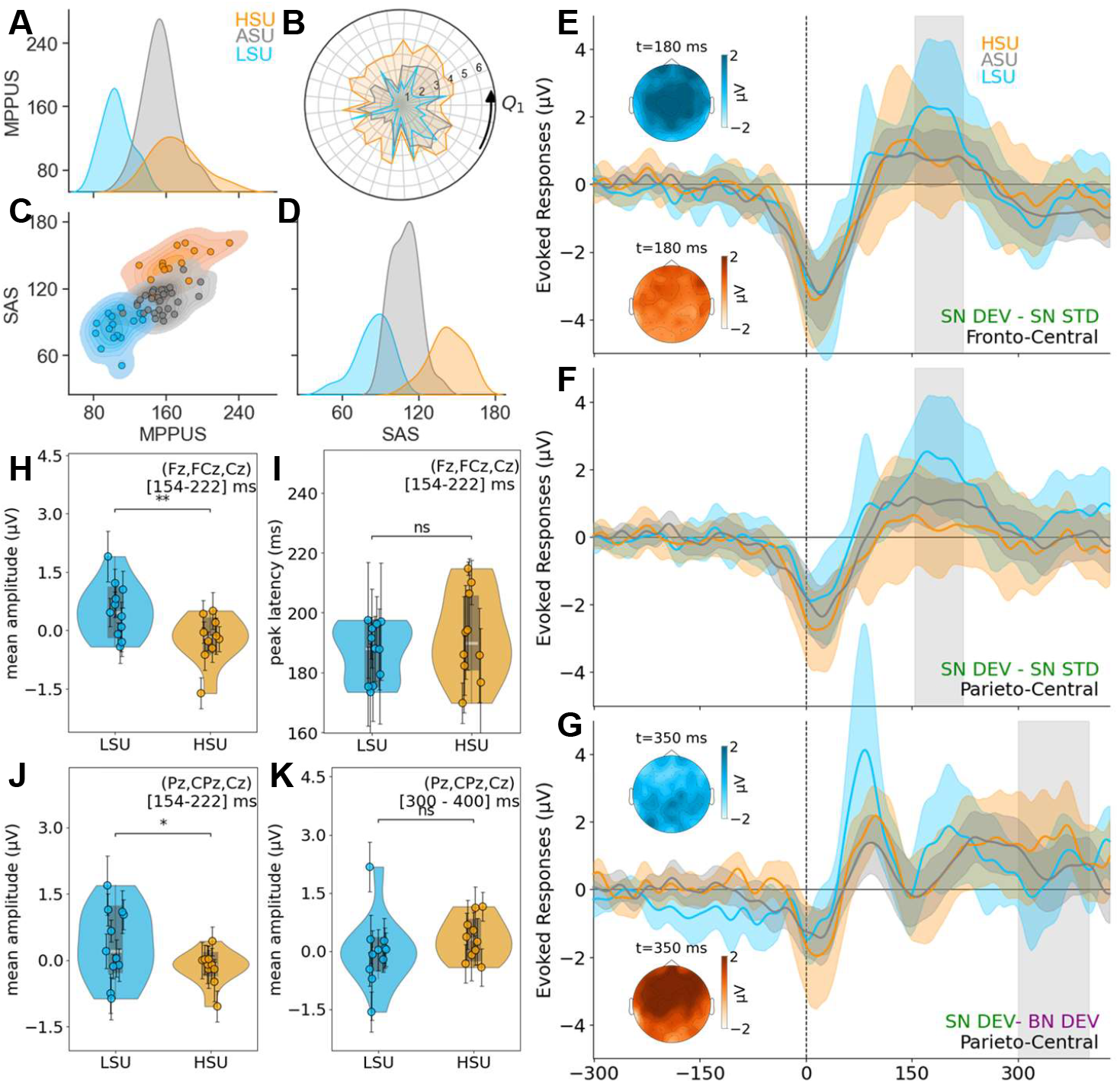
Differences in evoked responses across smartphone usage levels. **(A)** Distribution of Mobile Phone Problematic Use Scale (MPPUS) scores for HSU (orange), ASU (grey) and LSU (blue) groups. **(B)** Radar plot showing average SAS item-wise responses with a maximum possible score of 6 per question. **(C)** Joint density plot illustrating the relationship between SAS and MPPUS scores, highlighting clear separation among HSU, ASU and LSU participants. **(D)** Distribution of Smartphone Addiction Scale (SAS) scores across groups. **(E, F)** Grand-averaged N1-aligned SN DEV – SN STD at frontocentral (Cz, FCz, Fz) and parieto-central (Pz, CPz, Cz) locations. Topographic maps at 180 ms reveal spatial patterns consistent with the P3a ERP. **(G)** Grand-averaged N1-aligned SN DEV – BN DEV at parieto-central (Pz, CPz, Cz) location. Scalp topographies at 350 ms. **(E-G)** Shaded areas represent 95% confidence intervals. **(I)** Group differences in peak latency of P3a, **(H, J)** in average amplitude of P3a at the two electrode locations, and **(K)** in average amplitude of LPP for the deviant responses. **(H-K)** Violin plots show the distribution of individual participants’ ERP measures for the LSU, and HSU groups. White dots represent individual subject means, and black lines show within-subject standard deviations.

To assess whether data-driven groupings would reveal similar usage profiles, a spectral clustering analysis was conducted. Three clusters were identified, with group sizes of 14, 28, and 15 participants. Cluster quality metrics indicated moderately separable structure (Silhouette Score = 0.094; Calinski-Harabasz Score = 9.2; Davies-Bouldin Score = 2.36). These clusters were closely aligned with the average individual SAS item scores within HSU, LSU, and ASU categories (Fig. 4B). Of the total 57 participants, 14 were categorized as HSU, 15 as LSU, and 28 as ASU (Fig. 4D). Based on reliable identification of both N1 and P2 peaks, 42 participants were included in ERP analyses, comprising 10 HSU and 11 LSU individuals.

#### Comparable MMN Responses Across User Groups

Group-level comparisons of ERPs were performed between HSU and LSU for the SN (and BN) deviant response at the centro-frontal electrodes (Fig. 4E, S6). For the N1-aligned MMN window, no significant differences were observed between HSU and LSU in peak latency (0.0268 ± 0.0121 s vs. 0.0269 ± 0.0089 s; t ≈ 0.00, p = 0.998) or mean amplitude (-0.536 ± 0.391 μV vs. -0.730 ± 0.411 μV; t = 0.30, p = 0.772) for SN. These results indicate that early sensory deviance processing in the MMN range was comparable between heavy and light smartphone users.

#### Reduced Novelty P3a in HSU

In contrast, differences emerged in the later positive window corresponding to the P3a response in time window 154 ms - 222 ms from the N1 event onset (Fig. 4E shaded region). Mean amplitude differed significantly between the groups, with LSU showing a pronounced positive shift (0.523 ± 0.415 μV), whereas HSU did not (-0.224 ± 0.360 μV), yielding a significant contrast (t = -2.61, p = 0.0085, one-sided unpaired t-test) (Fig. 4H). No significant group differences were found for P3a peak latency (p = 0.81) (Fig 4I).

A corresponding analysis at the parieto-central electrodes, where the P3a typically reaches maximal amplitude (Fig. 4F), revealed a similar pattern: P3a peak latency did not differ between HSU and LSU (190.7 ± 11.7 ms vs. 183.7 ± 11.6 ms; t = 1.07, p = 0.85), while the mean amplitude indicated reduced positivity in HSU. LSU demonstrated a clear positive deflection (0.365 ± 0.406 μV), whereas HSU showed a negative amplitude (-0.164 ± 0.359 μV), resulting in a statistically significant group difference (t = -1.91, p = 0.037, one-sided unpaired t-test) (Fig. 4J).

#### Earlier Late Positive Potential (LPP) in HSU

Finally, we examined a later positive component over parieto-central sites corresponding to a delayed late positive potential (LPP). This response was quantified by subtracting the SN deviant waveform from the BN deviant waveform within the time window of 300 ms – 400 ms relative to the N1 event onset (Fig. 4G). For this component, peak latency differed significantly between groups: HSU exhibited an earlier LPP peak (349.79 ±14.5 ms) compared to LSU (362.95 ± 16.5 ms; t = -1.80, p = 0.044). This finding suggests a shift in the timing of later cognitive processing associated with higher smartphone use. In contrast, the mean amplitude of the LPP did not significantly differ between groups (HSU: 0.330 ± 0.525 μV; LSU: -0.011 ± 0.571 μV; t = 1.05, p = 0.847), indicating that the overall magnitude of this late parietal positivity was comparable across users. We do note, however, that the grand averaged HSU LPP remained consistently elevated while LSU showed large fluctuations.

## Discussion

The present study investigated how personalized smartphone notifications modulate different auditory ERPs, and how these responses differ with smartphone usage. Using a task-irrelevant auditory oddball paradigm with naturalistic auditory stimuli, we observed distinct neural signatures associated with personalized notification, suggesting that familiarity and usage intensity shape auditory processing at both early and late stages. One critical challenge with this ecological setup using personalized notifications is to compare neural responses across variable stimuli. A key feature of our analysis allowing us to overcome this challenge, was to use each subject’s auditory N1 peak in response to the standard stimulus as an alignment marker rather than stimulus onset; further backed by an across-block difference wave computation. Finally, behavioral quantification using smartphone addiction scales enabled us to discern usage-dependent modulation of the ERP components, providing further insights into the mechanistic interpretation of the observed effects across the ERP hierarchy.

Among the early auditory ERP components, we observed clearly dissociable adaptation and learning effects across standards and deviants. Both personalized (SN) and generic (BN) standard tones exhibited amplitude reductions from early to late trials, consistent with sensory adaptation to predictable stimuli ^31,32^. In contrast, no adaptation was observed for deviant responses, reflecting their continued violation of established auditory predictions. Notably, P2 amplitudes showed a distinct pattern: while P2 responses to standards remained stable across trials for both stimulus types, deviant BN tones exhibited a significant increase in P2 amplitude from early to late trials. Deviant SN tones, by contrast, elicited a positive P2 from the outset and showed no significant change over time. This pattern suggests that generic deviant tones undergo experience-dependent strengthening ^21^ of auditory object representations during the experiment, whereas personalized notification sounds engage an already well-established representation that does not require further learning^33^.

The mismatch negativity component provided clear evidence for stimulus-dependent modulation of pre-attentive auditory processing. Personalized notification sounds elicited MMNs that were both earlier and larger than those evoked by generic tones, indicating faster and stronger automatic prediction-error signaling, aligning with prior findings by ^29^. Notably, these MMN effects did not differ between high and low smartphone users. This dissociation suggests that personalized notifications engage a stable auditory representation shared across users, yielding robust pre-attentive salience detection that is largely independent of usage intensity^28,34–36^. Given the ubiquity of smartphones in contemporary environments, even individuals with relatively low usage are likely to possess well-established representations of notification sounds, potentially limiting the extent to which early predictive processes vary as a function of habitual exposure. Future studies spanning broader ranges of familiarity or exposure may help clarify the conditions under which usage-dependent modulation of early predictive processing emerges.

We did not observe a difference between SN and BN evoked P3a responses on average, although differences were observed between the LSU and HSU groups. While personalized notifications are more meaningful, they are also less novel than generic tones, and these opposing influences may plausibly counterbalance each other at the level of novelty-driven orienting reflected by the P3a. The pronounced group differences in P3a observed between low and high smartphone users may further reflect usage-dependent modulation of this balance: low-use individuals exhibited a clear positive P3a, consistent with intact novelty-related orienting, whereas high-use individuals showed attenuated or absent P3a responses, indicating reduced engagement of novelty-driven orienting mechanisms with increased habitual exposure.

Late positive potentials further revealed dissociable effects of stimulus identity and smartphone use on evaluative processing. Personalized notification sounds elicited a negative-going deflection in the LPP window compared to generic tones, which remained near baseline. This polarity difference suggests that personalized notifications engage a qualitatively distinct mode of late-stage processing, potentially reflecting reduced sustained evaluative engagement or task-driven active suppression of overlearned cues. In contrast, smartphone use modulated the temporal dynamics of the LPP rather than its magnitude: high smartphone users exhibited an earlier LPP peak compared to low users, despite comparable mean amplitudes. Taken together, these findings indicate that while stimulus personalization shapes the qualitative nature of late evaluative processing, habitual smartphone use influences the speed with which such processing is resolved. This pattern is consistent with a processing cascade in which personalized digital cues are rapidly detected, efficiently evaluated, and disengaged more quickly in individuals with greater habitual exposure.

A common assumption in discussions of digital distraction is that repeated exposure to personalized cues progressively amplifies their salience at early sensory stages, rendering habitual users increasingly reactive to notifications. The present findings place important constraints on this view. At a systems level, early auditory processing was primarily determined by stimulus identity and long-term familiarity, supporting robust and rapid detection of personalized cues that were shared across users and insensitive to usage intensity. In contrast, individual differences in smartphone use emerged only at later stages of processing associated with attentional orienting and extended evaluation. Together, this dissociation suggests that habitual exposure to digital signals is more likely to shape how salient information is handled downstream than how it is initially detected. This pattern highlights the importance of distinguishing early sensory salience from later attentional and evaluative processes when considering the cognitive impact of modern digital environments.

## Acknowledgements

The authors would like to acknowledge the insightful discussions and feedback from Dr. Vignesh Muralidharan, Center for Brain Science and Applications, Indian Institute of Technology, Jodhpur.

## Data availability

All anonymized data is available publicly in a clean, standardized BIDS format on the OpenNeuro repository at https://doi.org/10.18112/openneuro.ds007322.v1.0.0.

All experiment design and analysis code is available publicly at https://github.com/csndl-iitd/ecf-exp2-notif-mmn.

In an effort to promote open and replicable science, most study design and analysis decisions and unreported analysis directions (with corresponding discussions) are available publicly as issues in the above repository.

## Author contributions

SRG and PM designed research; PM performed experiments; SRG and PM analyzed data; SRG and PM wrote the paper; TKG provided equipment and supervision and project administration for PM.

