## Supplementary Information for "Personalized smartphone notifications bias auditory salience across processing stages"

**Appendix 1: Different Auditory Stimuli and their audio profile**

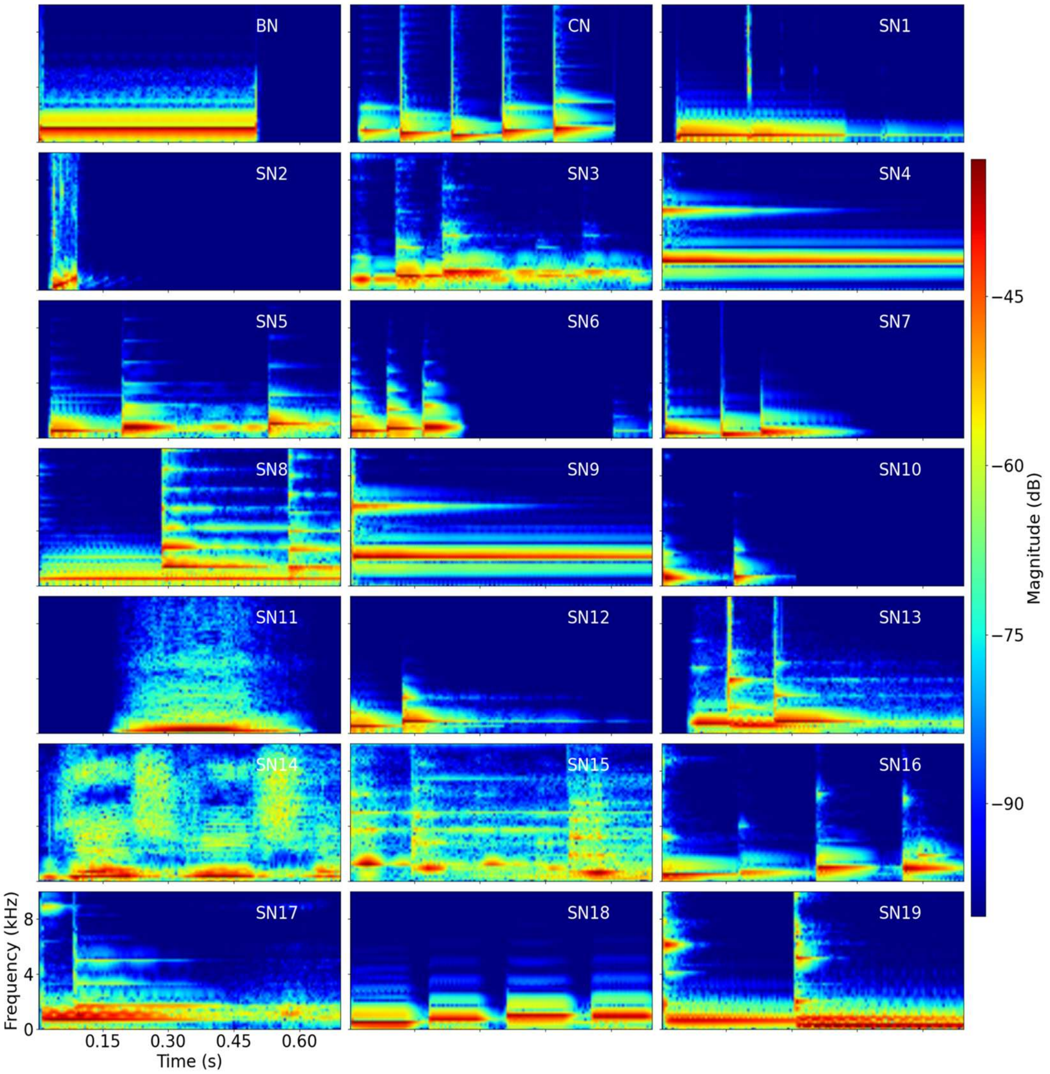


**Figure S1: Time–frequency representations (spectrograms) of background and smartphone notification sounds.** Each panel shows the spectral energy distribution (magnitude in dB) over time (s) and frequency (kHz) for the background noise (BN), control notification (CN), and individual smartphone notification sounds (SN1–SN19). Warmer colors indicate higher energy, illustrating the diversity in temporal structure and spectral content across notification types.

**Appendix 2: N1 – P2 SEM based subject selection and ERP alignment**

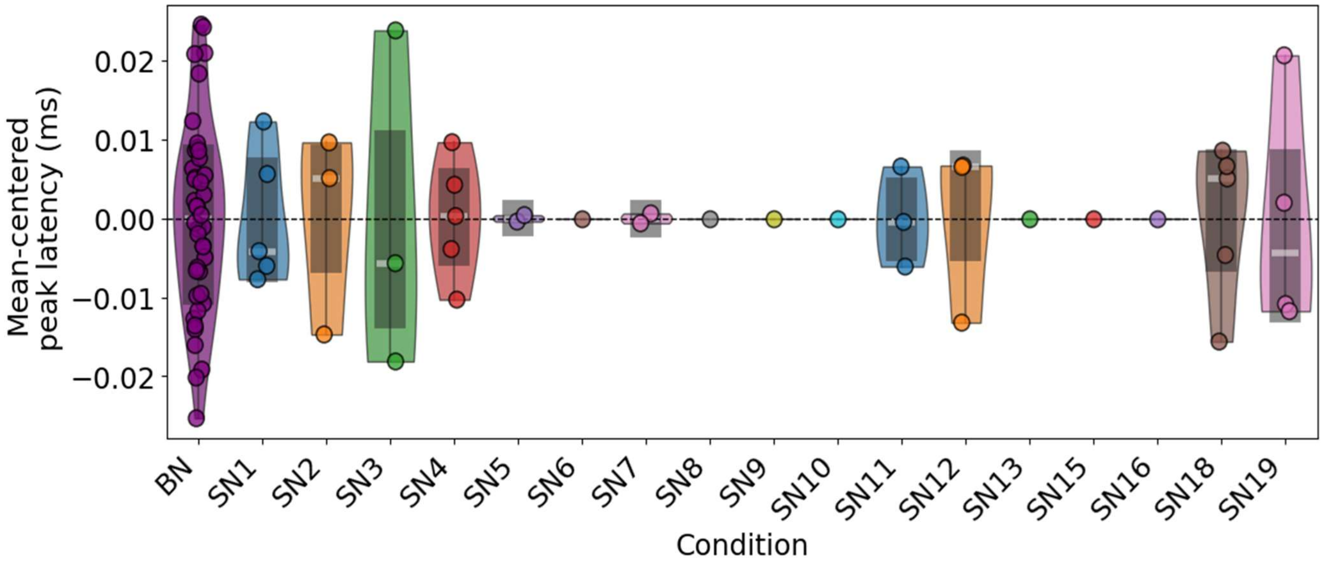


**Figure S2: Mean-centered peak N1 latency values for BN (purple) and SN (other colors)**. Each violin represents the distribution within a condition, with overlaid points showing individual subject means. Mean-centering ensures all conditions have a mean of zero, highlighting differences in variability of N1 peak latency. A total of 19 distinct personalized smartphone notifications were used in the study.

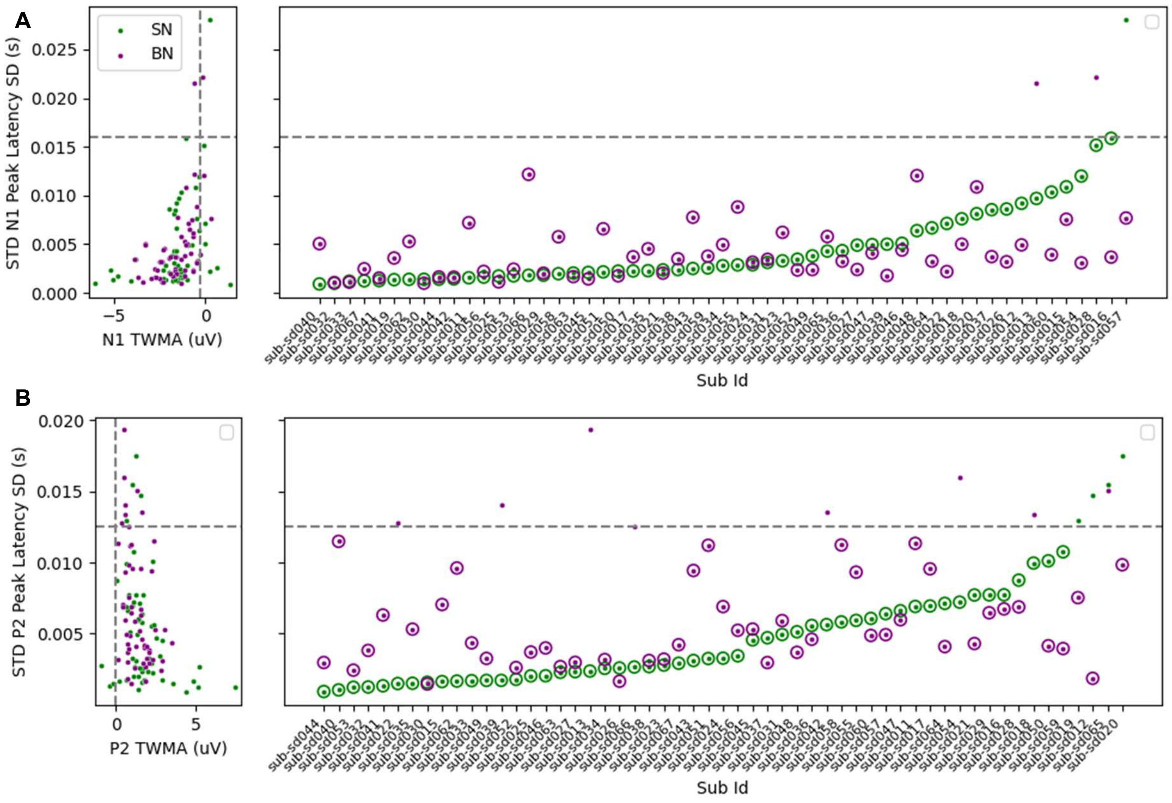


**Figure S3. Subject selection based on N1 and P2 ERP peak latency reliability for standard tones under smartphone notification (SN) and Beep notification (BN) conditions.** The y-axis represents the standard deviation (SD) of peak latency estimates for the corresponding ERP component (N1 or P2); higher SD values indicate poorer peak estimation reliability.

(A) Individual subjects' N1 peak latency SDs.

(B) Individual subjects' P2 peak latency SDs.

Subjects with SD values below the dotted threshold lines (N1: 0.016 s; P2: 0.012 s) for both SN and BN conditions were included in subsequent analyses.

**Appendix 3: Smartphone usage related questionnaires**

**3.1. Smartphone addiction scale (SAS)**

Participants indicated their responses on a six-point Likert-type scale ranging from 1 (*strongly disagree*) to 6 (*strongly agree*).

1. Missing planned work due to smartphone use.
2. Having a hard time concentrating in class, while doing assignments, or while working due to smartphone use.
3. Experiencing lightheadedness or blurred vision due to excessive smartphone use.
4. Feeling pain in the wrists or at the back of the neck while using a smartphone.
5. Feeling tired and lacking adequate sleep due to excessive smartphone use.
6. Feeling calm or cozy while using a smartphone.
7. Feeling pleasant or excited while using a smartphone.
8. Feeling confident while using a smartphone.
9. Being able to get rid of stress with a smartphone.
10. There is nothing more fun to do than using my smartphone.
11. My life would be empty without my smartphone.
12. Feeling most liberal while using a smartphone.
13. Using a smartphone is the most fun thing to do.
14. Won't be able to stand not having a smartphone.
15. Feeling impatient and fretful when I am not holding my smartphone.
16. Having my smartphone in my mind even when I am not using it.
17. I will never give up using my smartphone even when my daily life is already greatly affected by it.
18. Getting irritated when bothered while using my smartphone.
19. Bringing my smartphone to the toilet even when I am in a hurry to get there.
20. Feeling great meeting more people via smartphone use.
21. Feeling that my relationships with my smartphone buddies are more intimate than my relationships with my real-life friends.
22. Not being able to use my smartphone would be as painful as losing a friend.
23. Feeling that my smartphone buddies understand me better than my real-life friends.
24. Constantly checking my smartphone so as not to miss conversations between other people on Twitter or Facebook.
25. Checking SNS (Social Networking Service) sites like Twitter or Facebook right after waking up.
26. Preferring talking with my smartphone buddies to hanging out with my real-life friends or with the other members of my family.
27. Preferring searching from my smartphone to asking other people.
28. My fully charged battery does not last for one whole day.
29. Using my smartphone longer than I had intended.
30. Feeling the urge to use my smartphone again right after I stopped using it.
31. Having tried time and again to shorten my smartphone use time, but failing all the time.
32. Always thinking that I should shorten my smartphone use time.
33. The people around me tell me that I use my smartphone too much.

**3.2. Mobile Phone Problematic Use Scale (MPPUS)**

Participants indicated their responses on a ten-point Likert-type scale ranging from 1 (*not true*) to 10 (*Extremely true*)

1. I can never spend enough time on my mobile phone.

2. I have used my mobile phone to make myself feel better when I was feeling down.
3. I find myself occupied on my mobile phone when I should be doing other things, and it causes problems.
4. All my friends own a mobile phone.
5. I have tried to hide from others how much time I spend on my mobile phone.
6. I lose sleep due to the time I spend on my mobile phone.
7. I have received mobile phone bills I could not afford to pay.
8. When out of range for some time, I become preoccupied with the thought of missing a call.
9. Sometimes, when I am on the mobile phone and I am doing other things, I get carried away with the conversation and I don’t pay attention to what I am doing.
10. The time I spend on the mobile phone has increased over the last 12 months.
11. I have used my mobile phone to talk to others when I was feeling isolated.
12. I have attempted to spend less time on my mobile phone but am unable to.
13. I find it difficult to switch off my mobile phone.
14. I feel anxious if I have not checked for messages or switched on my mobile phone for some time.
15. I have frequent dreams about the mobile phone.
16. My friends and family complain about my use of the mobile phone.
17. If I don't have a mobile phone, my friends would find it hard to get in touch with me.
18. My productivity has decreased as a direct result of the time I spend on the mobile phone.
19. I have aches and pains that are associated with my mobile phone use.
20. I find myself engaged on the mobile phone for longer periods of time than intended.
21. There are times when I would rather use the mobile phone than deal with other more pressing issues.
22. I am often late for appointments because I'm engaged on the mobile phone when I shouldn't be.
23. I become irritable if I have to switch off my mobile phone for meetings, dinner engagements, or at the movies.
24. I have been told that I spend too much time on my mobile phone.
25. More than once I have been in trouble because my mobile phone has gone off during a meeting, lecture, or in a theatre.
26. My friends don't like it when my mobile phone is switched off.
27. I feel lost without my mobile phone.

**3.3. Additional Questions about smartphone usages pattern**

Q1. How much do you agree with the statement: I am addicted to my smartphone?

- Strongly disagree
- Disagree
- Neutral
- Agree
- Strongly agree

Q2. My phone is on

Indicate your answers specifically for the app that you get most notifications on.

| | Never | Sometimes | Often | Always |
|---|---|---|---|---|
| Silent/DND | ☐ | ☐ | ☐ | ☐ |
| Vibrate only | ☐ | ☐ | ☐ | ☐ |

| | | | | |
|---|---|---|---|---|
| loud | ☐ | ☐ | ☐ | ☐ |

Q3. Do you receive notifications on some other device instead (like a smartwatch)?

Yes No

Q4. What emotions are typically associated with receiving smartphone notifications?

- Annoying
- Sad
- Neutral
- Exciting
- Happy

Q5. How did you feel about the notifications during the first half of the experiment?

| Not Annoying | 1 | 2 | 3 | 4 | 5 | Extremely Annoying |
|---|---|---|---|---|---|---|
| | ☐ | ☐ | ☐ | ☐ | ☐ | |

Q6. How did you feel about the notifications during the 2nd half of the experiment?

| Not Annoying | 1 | 2 | 3 | 4 | 5 | Extremely Annoying |
|---|---|---|---|---|---|---|
| | ☐ | ☐ | ☐ | ☐ | ☐ | |

**Appendix 4: Within block analyses**

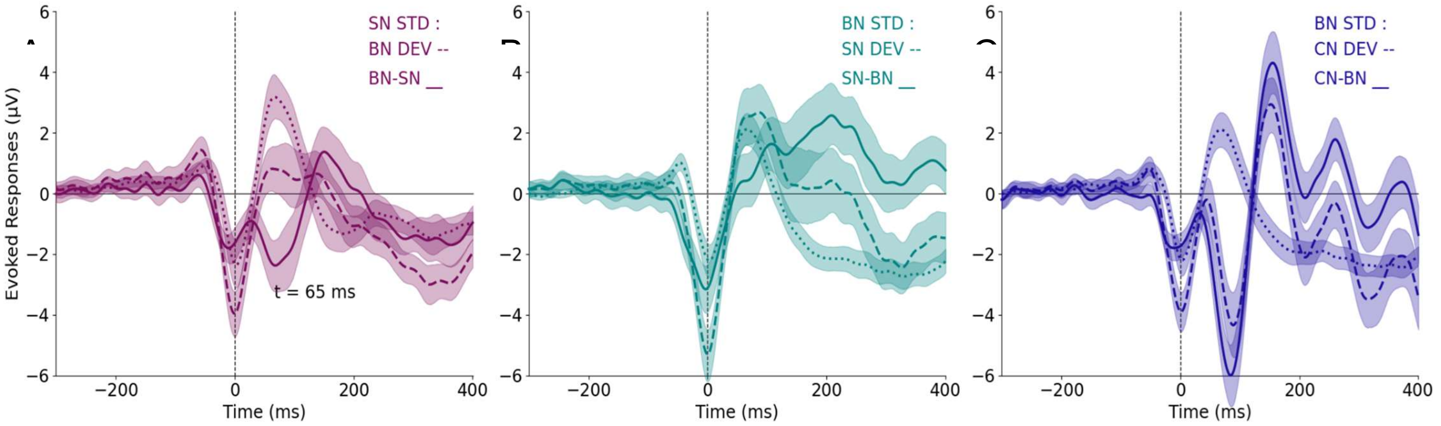


**Figure S4.** Within-block analysis of responses to standard (STD) and deviant (DEV) tones. The dotted line represents the response to standard stimuli, the dashed line represents the response to deviant stimuli within the same block, and the solid line represents the difference between STD and DEV responses within that block.(A) Block with SN as STD and BN as DEV. (B) Block with SN as both STD and DEV. (C) Block with BN as STD and CN as DEV within that block.

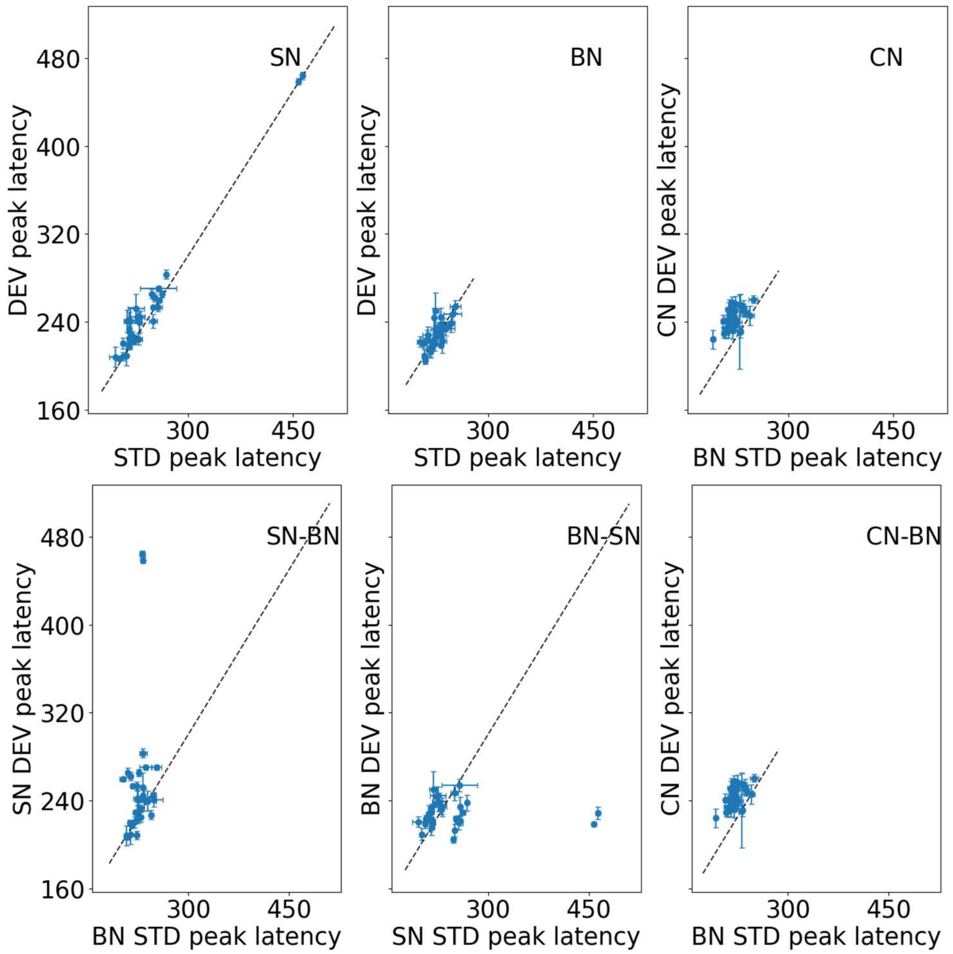


**Figure S5. P**eak latency of N1 in response to the same tone as deviant (DEV) and standard (STD) (top row); in response to respective deviant and standard tones within the same block (bottom row).

**Appendix 5 - BN MMN response for the HSU, LSU, ASU at fronto-central and parieto-central locations**

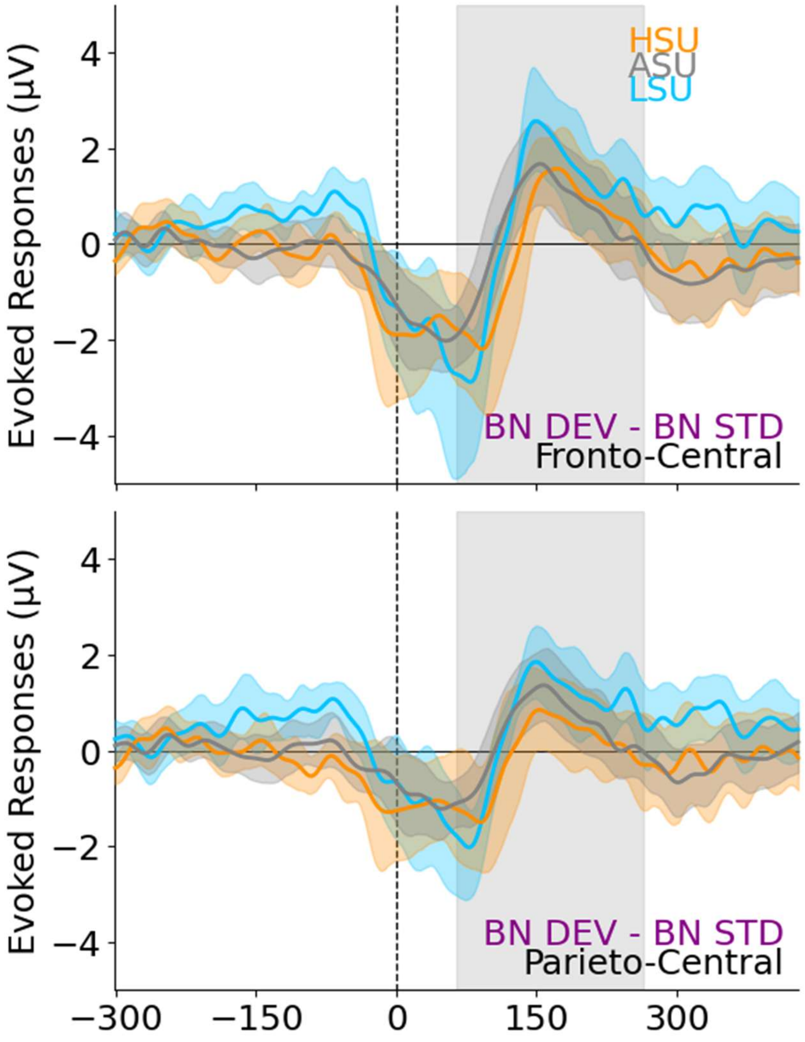


**Figure S6.** HSU, ASU and LSU MMN for Fronto central and parieto-central electrodes
